# Protein CoAlation on TXNRD2 regulates mitochondrial thioredoxin system to protect against ferroptosis

**DOI:** 10.1101/2024.05.16.594391

**Authors:** Chao-Chieh Lin, Yi-Tzu Lin, Ssu-Yu Chen, Yasaman Setayeshpour, Yubin Chen, Denise E Dunn, Erik J Soderblom, Guo-Fang Zhang, Valeriy Filonenko, Suh Young Jeong, Scott Floyd, Susan J Hayflick, Ivan Gout, Jen-Tsan Chi

## Abstract

The Cystine-xCT transporter-Glutathione (GSH)-GPX4 axis is the canonical pathway to protect against ferroptosis. While not required for ferroptosis-inducing compounds (FINs) targeting GPX4, FINs targeting the xCT transporter require mitochondria and its lipid peroxidation to trigger ferroptosis. However, the mechanism underlying the difference between these FINs is still unknown. Given that cysteine is also required for coenzyme A (CoA) biosynthesis, here we show that CoA supplementation specifically prevents ferroptosis induced by xCT inhibitors but not GPX4 inhibitors. We find that, auranofin, a thioredoxin reductase inhibitor, abolishes the protective effect of CoA. We also find that CoA availability determines the enzymatic activity of thioredoxin reductase, but not thioredoxin. Importantly, the mitochondrial thioredoxin system, but not the cytosolic thioredoxin system, determines CoA-mediated ferroptosis inhibition. Our data show that the CoA regulates the *in vitro* enzymatic activity of mitochondrial thioredoxin reductase (TXNRD2) by covalently modifying the thiol group of cysteine (CoAlation) on Cys-483. Replacing Cys-483 with alanine on TXNRD2 abolishes its *in vitro* enzymatic activity and ability to protect cells from ferroptosis. Targeting xCT to limit cysteine import and, therefore, CoA biosynthesis reduced CoAlation on TXNRD2, an effect that was rescued by CoA supplementation. Furthermore, the fibroblasts from patients with disrupted CoA metabolism demonstrate increased mitochondrial lipid peroxidation. In organotypic brain slice cultures, inhibition of CoA biosynthesis leads to an oxidized thioredoxin system, mitochondrial lipid peroxidation, and loss in cell viability, which were all rescued by ferrostatin-1. These findings identify CoA-mediated post-translation modification to regulate the thioredoxin system as an alternative ferroptosis protection pathway with potential clinical relevance for patients with disrupted CoA metabolism.

## Introduction

The *de novo* biosynthesis of coenzyme A (CoA) requires the phosphorylation of extracellular pantothenate (Vitamin B5) by pantothenate kinase (PANK), followed by the subsequent incorporation of cysteine and ATP^1^. Gene mutations in two of the genes involved in CoA biosynthesis, including mitochondrial pantothenate kinase 2 (*PANK2*) and CoA synthase (*COASY*) lead to PKAN (Pantothenate Kinase-Associated Neurodegeneration) and CoPAN (COASY Protein-Associated Neurodegeneration), respectively^2^. Both PKAN and CoPAN belong to a group of inherited neurological disorders termed neurodegeneration with brain iron accumulation (NBIA). These disorders are characterized by progressive dystonia, dysarthria, spasticity, parkinsonism, and other neuropsychiatric abnormalities. However, there is currently no cure for PKAN and CoPAN, and the pathogenesis mechanisms remains unclear^3^

CoA and its thioester derivatives are involved in diverse functions, including protein acetylation, Krebs cycle, amino acid metabolism, fatty acid synthesis, and regulation of gene expression^4^. While the cytosolic concentrations of CoA were estimated to be at micromolar levels, mitochondrial CoA concentrations are significantly higher at millimolar levels, which may imply that the functional role of CoA is more crucial in mitochondria^5^. Recently, a novel function of CoA has been discovered involving the covalent attachment of CoA to the thiol group of cysteine residues in target proteins under oxidative stress, and termed CoAlation^6^. More than 2000 mammalian and bacterial proteins have been identified to be CoAlated^6^. However, much remains unknown about its regulation and biological functions^7^.

In pancreatic tumors, CoA has been proposed to increase coenzyme Q_10_ via the mevalonate pathway to protect cancer cells against ferroptosis^8^. Ferroptosis is a newly appreciated death mechanism characterized by iron dependency, oxidative stress, and lipid peroxidation. Ferroptosis can be triggered by various ferroptosis-inducing compounds (FINs), which can be categorized into different classes based on their targets and mechanisms. The class I FINs, including erastin and sulfasalazine (SAS), target xCT transporter to block cystine uptake. Class II FINs, such as RSL3 and ML162, target the enzymatic activity of GPX4. Class III FINs deplete both GPX4 protein and ubiquinone^9^. Notably, although both class I and class II FINs act upon the same Cystine-xCT-glutathione (GSH)-GPX4 axis, only class I, but not class II, FINs induce mitochondrial lipid peroxidation^10,11^. While the necessity of mitochondria and its lipid peroxidation for class 1 FINs has been established^10^, the precise mechanism remains elusive.

The thioredoxin system works in parallel with the glutathione system to regulate the cellular redox environment and protect against ferroptosis^12^. The thioredoxin system facilitates the sequential transfer of electrons from NADPH to thioredoxin reductase (TXNRD), then to thioredoxin (TXN), and finally to peroxiredoxin (PRDX), enabling the elimination of intracellular reactive oxygen species (ROS). Buthionine sulfoximine (BSO) inhibits GSH biosynthesis by targeting gamma-glutamylcysteine synthetase. However, BSO is a far less potent FIN than erastin and other FINs. One study suggests that inhibiting GSH with BSO increases intracellular cystine levels, subsequently enhancing thioredoxin system to prevent ferroptosis^13^. Therefore, combining BSO with thioredoxin reductase inhibitor (auranofin) can synergistically induce cell death^13^. Nevertheless, much remains unknown about the mechanism by which elevated cystine levels regulate the thioredoxin system.

In this study, we found that CoA is a class I FIN-specific inhibitor of ferroptosis. The availability of CoA modulates the redox status of mitochondrial thioredoxin system to prevent mitochondrial lipid peroxidation and protect against ferroptosis. Mechanistically, CoA covalently modifies (CoAlation) Cys-483 of mitochondrial thioredoxin reductase (TXNRD2) to enhance its enzymatic activity and prevent mitochondrial lipid peroxidation during ferroptosis. Consistently, fibroblasts from PKAN patients also show an increase in mitochondrial lipid peroxidation. Importantly, these findings were validated in organotypic brain slice cultures (OBSC). Our findings elucidate class I FIN-specific mitochondrial lipid peroxidation and the requirement of cysteine incorporation for CoA to regulate the thioredoxin system, with potential clinical relevance for both PKAN and CoPAN patients.

## Results

### CoA-mediated ferroptosis inhibition is partially regulated by GSH

Supplementation of CoA in the culture medium has been shown to protect mouse embryonic fibroblasts and pancreatic tumors from ferroptosis^8,14^. Extracellular CoA was proposed to be first degraded to 4’-phosphopantetheine (4’-PPT), which can enter cells as an alternative CoA biosynthetic pathway^15^. To confirm that supplementing culture media with CoA indeed increases intracellular CoA levels in our model, we treated HT-1080 cells with two different concentrations of CoA for 18 hours and measured the intracellular levels of CoA and acetyl-CoA by mass spectrometry (Fig. 1A-B). We confirmed that the CoA supplementation significantly increased intracellular CoA levels (Fig. 1A). The increase in acetyl-CoA levels further confirmed that the supplemented CoA can serve as an acetyl group carrier (Fig. 1B). Next, we measured the protective capacity of CoA and several canonical ferroptosis inhibitors, including deferoxamine (DFO), ferrostatin-1 (Fer-1), liproxstatin-1 (lipro), and Trolox, against erastin-induced ferroptosis in HT-1080 cells. We found that the ferroptosis protective effect of CoA was comparable to those of the canonical ferroptosis inhibitors (Fig. 1C). Subsequently, we confirmed that the ferroptosis inhibitory effect of CoA could be reproduced in five different cancer cell lines (HEK-293, 786-O, H1975, RCC4, and MDA-MB-231 cells, Supplemental Fig. 1A-E). Consistently, CoA also abolished other molecular features of erastin-induced ferroptosis, including lipid peroxidation (Fig. 1D-E) and membrane rupture (Fig. 1F-G). Moreover, we found that CoA only inhibited ferroptosis triggered by class I FINs (erastin, BSO, and sulfasalazine) (Fig. 1C, Supplemental Fig. 1F-G), but not class II FINs (ML162 and RSL3) (Fig. 1H-I) or class III FIN (FIN56) (Fig. 1J). Taken together, these results indicate that CoA-mediated inhibition of ferroptosis may regulate processes upstream of GPX4.

**Fig. 1.**
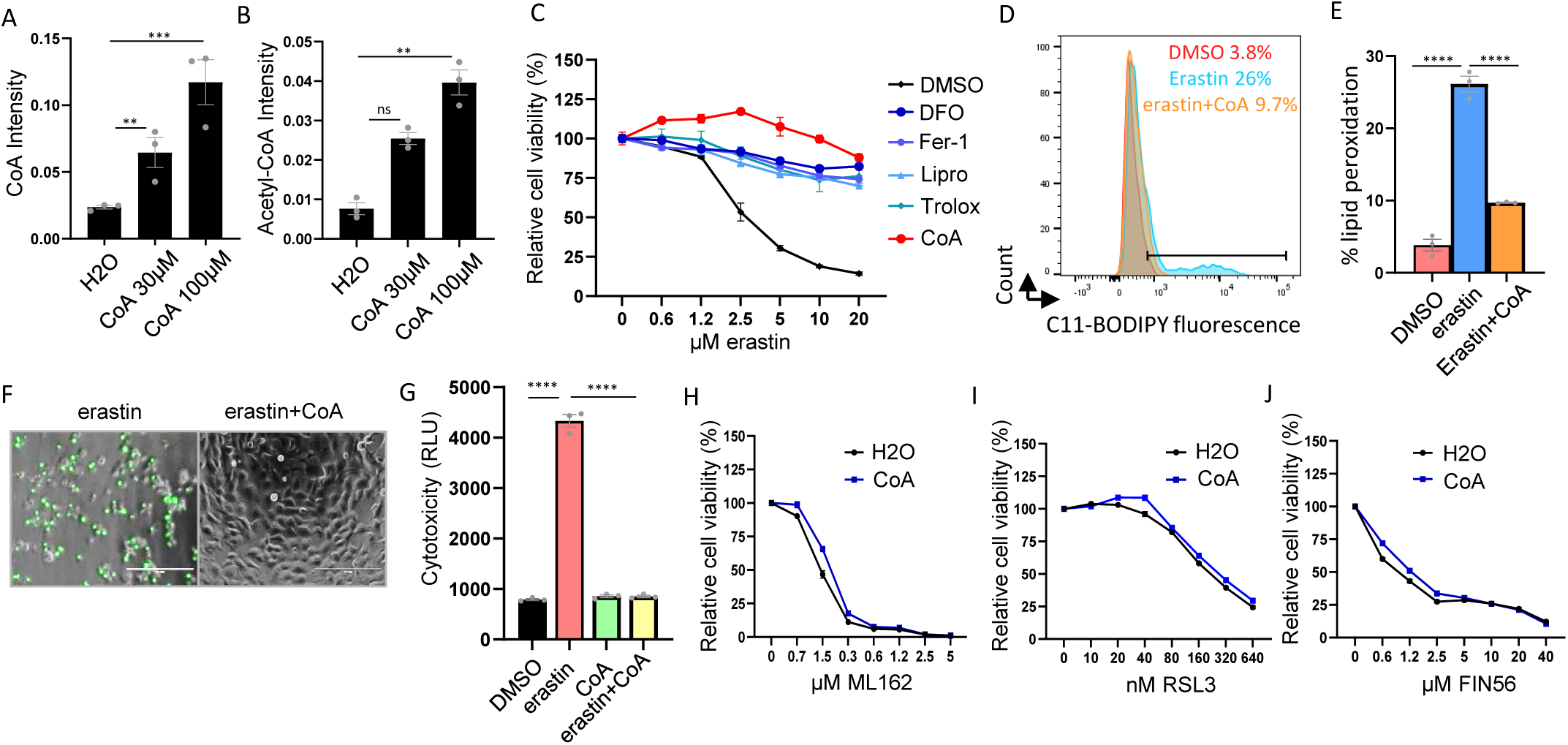
CoA is a class 1 FIN-specific ferroptosis inhibitor. (**A-B**) CoA supplementation increased the levels of intracellular CoA (**A**) and acetyl-CoA (**B**) as quantified by LC-MS/MS analysis. HT-1080 cells were treated with H2O, CoA (30 μM, 100 μM) for 18 hours for LC-MS/MS analysis. (**C**) CoA inhibited erastin-induced ferroptosis. HT-1080 cells were treated with increasing doses of erastin, either alone or in combination with CoA (100 μM) or deferoxamine (DFO, 80 μM), ferrostatin-1 (Fer-1, 10 μM), liproxstatin-1 (lipro, 2 μM), Trolox (100 μM). The cell viability was quantified by Cell-Titer Glo assay. (**D-E**) Erastin-induced lipid peroxidation (2 µM, 18 hours) in HT-1080 cells was inhibited by CoA treatment as determined by C11-BODIPY staining (**D**) and the quantification of % lipid peroxidation positive cells (**E**). (**F-G**) CoA (100 μM) inhibited erastin (2.5 μM, 20 hours)-induced membrane rupture in HT-1080 cells as observed by CellTox Green under fluorescence microscope (**F**) and quantified by a plate reader (**G**). (**H-I**) CoA (100 μM) failed to inhibit class 2 FIN-induced ferroptosis in HT-1080 cells including ML162 (20 hours) (**H**) and RSL3 (20 hours) (**I**). (**J**) CoA (100 μM) failed to inhibit class 3 FIN (FIN56)-induced ferroptosis in HT-1080 cells.

### The mitochondrial thioredoxin system determines CoA-mediated ferroptosis inhibition

To uncover other potential mechanisms of CoA-mediated ferroptosis inhibition, we performed a target compound screen to identify compounds that could abolish CoA-mediated ferroptosis protection. In pancreatic tumors, CoA was proposed to inhibit lipid peroxidation by synthesizing coenzyme Q10 for FSP1 via the mevalonate pathway ^8^. Thus, we determined whether the CoA-protected ferroptosis could be abolished by FSP1 inhibitor: FSEN1 (Supplemental Fig. 2A). While FSEN1 was able to sensitize ferroptosis in the control cells, FSEN1 did not abolish CoA-mediated inhibition of ferroptosis in HT-1080 cells (Supplemental Fig. 2A). Next, we evaluated the efficacy of various inhibitors targeting pathways recognized to be critical for ferroptosis protection to determine their potential to abolish CoA-mediated ferroptosis protection. These inhibitors included brequinar and teriflunomide^11^ (DHO dehydrogenase inhibitors, Supplemental Fig. 2B-C), Etomoxir^16^ (β-oxidation inhibitor, Supplemental Fig. 2D), methotrexate^17^ (dihydrofolate reductase inhibitor, Supplemental Fig. 2E), compound C^18^ (AMPK inhibitor, Supplemental Fig. 2F), TOFA^19^ (ACC inhibitor, Supplemental Fig. 2G), the inhibitors of the S-adenosyl homocysteine hydrolase of the trans-sulfuration pathway^20^ (Supplemental Fig. 2H). However, none of these compounds mitigated the ferroptosis protection effect of CoA (Supplemental Fig. 2A-H). Given that ACSL3 mediates the incorporation of CoA into monounsaturated fatty acids (MUFA)^21,22^, we conducted a genetic knockdown of ACSL3 and found that it had no effect on CoA-mediated protection against ferroptosis (Supplemental Fig. 2J).

Among the compounds tested, we found that both auranofin^13^ (thioredoxin reductase inhibitor) and ferroptocide^23^ (thioredoxin inhibitor) abolished CoA-mediated inhibition of ferroptosis (Fig. 2A, Supplemental Fig. 2K). Furthermore, CoA was unable to rescue auranofin or ferroptocide-induced cell death (Supplemental Fig. 2L-M). These data suggest that the thioredoxin system is critical in the CoA-mediated inhibition of ferroptosis. While any screens may have false negatives and our results did not completely rule out the involvement of other pathways, the ability of auranofin and ferroptocide prompted us to further investigate the thioredoxin pathways. Thus, we examined the thioredoxin and thioredoxin reductase activities in HT-1080 cell lysates when treated with erastin, CoA, and their combination (Fig. 2B, Supplemental Fig. 2N). While thioredoxin activity in HT-1080 cell lysates was not altered by erastin or CoA (Supplemental Fig. 2N), erastin significantly reduced thioredoxin reductase activity which was restored by CoA supplementation (Fig. 2B). However, the protein levels of both cytosolic and mitochondrial thioredoxin reductases (TXNRD1 and TXNRD2) were not affected by erastin or CoA (Supplemental Fig. 2O). Since erastin was known to reduce the intracellular levels of CoA, we speculated that reduced CoA levels may determine thioredoxin reductase activity during ferroptosis.

**Fig. 2.**
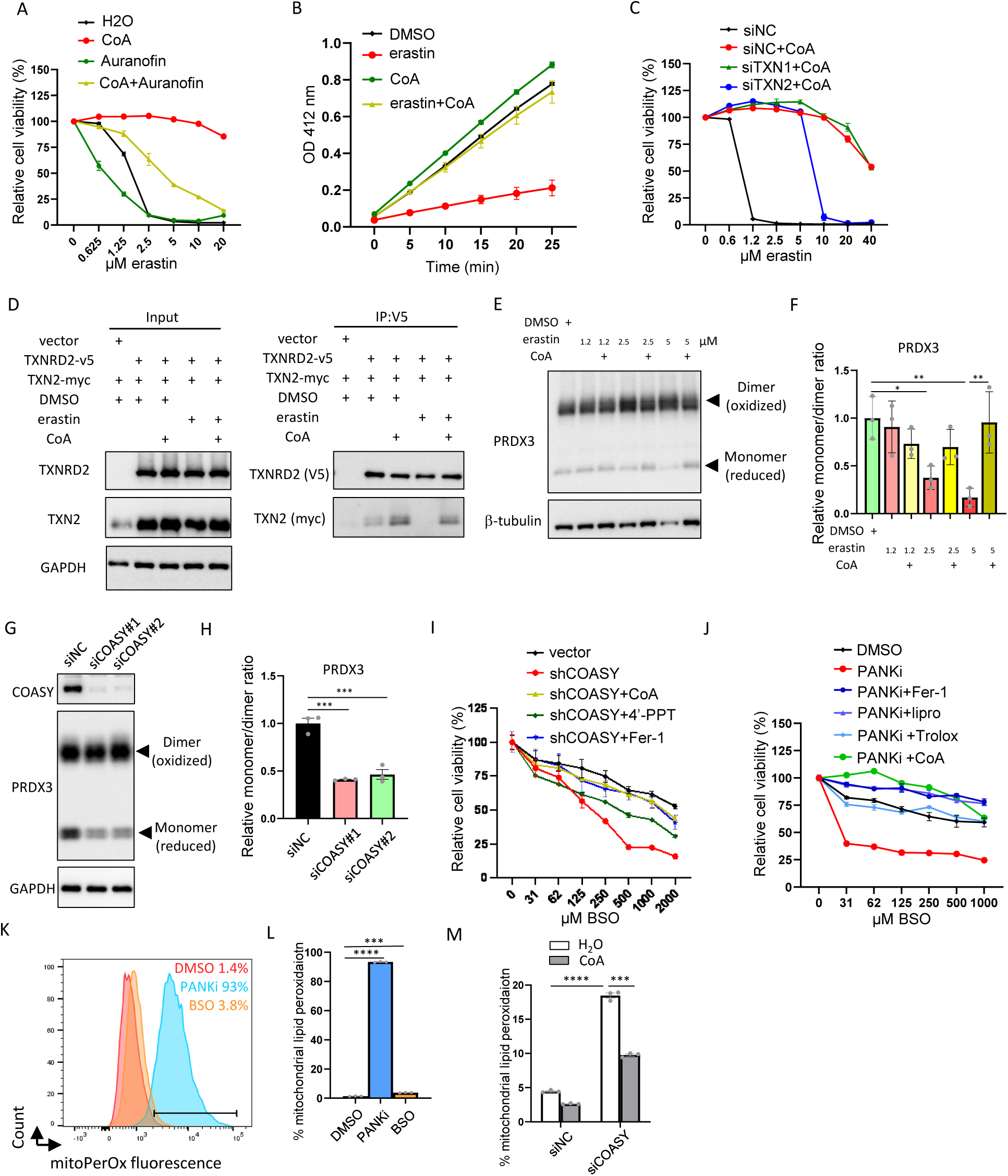
CoA regulates mitochondrial thioredoxin system. (**A**) Auranofin (0.5 μM) abolished CoA (100 μM)-mediated ferroptosis protection of HT-1080 cells caused by indicated levels of erastin for 20 hours as quantified by Cell-Titer Glo assay. (**B**) Erastin (1.25 μM, 16 hours) significantly repressed the thioredoxin reductase activity in HT-1080 cell lysates, which can be restored by CoA (100 μM) supplementation. (**C**) The knockdown of TXN2, but not TXN1, mitigated CoA-mediated ferroptosis protection. (**D**) The TXNRD2-TXN2 interaction was abolished by erastin, which can be restored by CoA supplement. HT-1080 cells overexpressing TXNRD2 and TXN2 were treated with erastin (2.5 μM, 18 hours) or in combination with CoA (100 μM) and were lysed with NEM for co-immunoprecipitation. (**E**) Erastin-induced decrease in the monomers (reduced and active forms) of PRDX3 in HT-1080 cells was rescued by CoA supplementation as determined by Western blots. (**F**) Quantification of the monomer/dimer ratio of PRDX3 upon erastin treatment with or without CoA supplementation. (**G**) COASY knockdown in HT-1080 cells by two independent COASY-targeting siRNAs reduced the monomer (reduced and active forms) of PRDX3 in Western blots. (**H**) Quantification of the monomer/dimer ratio of PRDX3 upon the knockdown of COASY. (**I**) COASY knockdown by COASY shRNA sensitized HT-1080 cells to BSO treatment, which was rescued by CoA (100 μM), 4’-phosphopantetheine (4’-PPT, 100 μM), ferrostatin-1 (Fer-1, 10 μM). (**J**) CoA inhibited ferroptosis induced by the combination of BSO and PANKi. HT-1080 cells were treated with PANKi (2.5 µM) and an increasing dose of BSO in combination with ferrostatin-1 (Fer-1, 10 μM), liproxstatin-1 (lipro, 2 μM), Trolox (100 μM), or CoA (100 μM). The cell viability was quantified by Cell-Titer Glo assay. (**K-L**) PANKi, but not BSO, increased mitochondrial lipid peroxidation (PANKi 2.5 µM, BSO 1 mM, 18 hours) in HT-1080 cells as determined by the sensor of mitochondrial lipid peroxidation (mitoPerOx) staining (**K**) and the quantification of % mitochondrial lipid peroxidation positive cells (**L**). (**M**) COASY knockdown in HT-1080 cells triggered mitochondrial lipid peroxidation, which was rescued by CoA treatment, as quantification by % mitochondrial lipid peroxidation (mitoPerOX) positive cells.

To determine the relative importance of cytosolic versus mitochondrial thioredoxin systems for CoA-mediated inhibition of ferroptosis, we employed pooled siRNA against cytosolic thioredoxin (TXN1) and mitochondrial thioredoxin (TXN2) to knockdown these two genes individually (Supplemental Fig. 2P). We found that the knockdown of TXN2, but not TXN1, abolished CoA-mediated ferroptosis inhibition (Fig. 2C), and this result was further validated using individual TXN2 siRNAs (Supplemental Fig. 2Q). These data indicate that mitochondrial thioredoxin system is required for CoA-mediated protection from ferroptosis.

While previous findings suggest that unutilized cysteine directly fuels the thioredoxin system to prevent BSO-induced ferroptosis^13^, we hypothesized that unutilized cysteine is converted to CoA to regulate ferroptosis. The thioredoxin system transfers electrons in the order of NADPH, TXNRD, TXN, and PRDXs to neutralize intracellular ROS. Given the importance of mitochondrial TXN2 (Fig. 2C), we focused on the mitochondrial thioredoxin system and examined the protein-protein interaction between TXNRD2 and its substrate TXN2. Through co-immunoprecipitation, we found that erastin treatment abolished the interaction between TXNRD2 and TXN2, while CoA supplementation rescued this interaction (Fig. 2D). The redox status of the thioredoxin system can be monitored by the monomer (reduced and active)/ dimer (oxidized and inactive) ratio of PRDXs^24^. Thus, we treated HT-1080 cells with erastin alone or combined with CoA to measure the monomer/dimer ratio of PRDX3 (Fig. 2E-F). We noticed that erastin decreased the monomer (reduced and active form) of mitochondrial PRDX3 (Fig. 2E-F), which were rescued by CoA supplementation (Fig. 2E-F). Our previous findings showed that the knockdown of CoA synthase (COASY) reduced intracellular CoA levels^25^. Therefore, we reduced CoA levels by knocking down COASY and assessed the redox status of the mitochondrial thioredoxin system by measuring the monomer/dimer ratio of PRDX3 (Fig. 2G). Consistently, COASY knockdown decreased the reduced form (monomer) of PRDX3 (Fig. 2H). This reduced monomer/dimer ratio suggests a defect in mitochondrial thioredoxin function upon erastin treatment (Fig. 2E, F) and COASY knockdown (Fig. 2G-H).

These results prompt us to hypothesize that GSH and CoA may mediate parallel pathways of ferroptosis protection. To determine the contribution of the erastin-depleted GSH and CoA on ferroptosis, we combined BSO (GSH inhibitor) and COASY knockdown (Fig. 2I). While BSO itself is known to be a much less potent inducer of ferroptosis, we found that combining BSO with COASY knockdown significantly decreased cell viability over extended incubation, which was rescued by supplementing CoA, 4’-PPT or ferrostatin-1 (Fig. 2I, Supplemental Fig. 2R). Similarly, a previous report showed that combining BSO (GSH inhibitor) and PANKi^26^ (CoA biosynthesis inhibitor) also triggers ferroptosis in pancreatic tumors^8^. Therefore, we combined BSO and PANKi and found such a combination led to dramatic cell death of HT-1080 cells that were rescued by CoA to a similar degree as ferroptosis inhibitors (ferrostatin-1, Liproxstatin-1 and Trolox) (Fig. 2J). Given that COASY knockdown led to impaired redox status of the mitochondrial thioredoxin system and the relevance of the mitochondrial thioredoxin system for CoA-mediated ferroptosis inhibition (Fig. 2C, G), we treated HT-1080 cells with PANKi or BSO. We found that inhibition of CoA by PANKi, compared with BSO, indeed led to robust mitochondrial lipid peroxidation (Fig. 2K-L). Consistently, genetic inhibition of CoA synthesis by COASY siRNA significantly increased mitochondrial lipid peroxidation, which could be abolished by CoA supplementation (Fig. 2M). Taken together, while previous findings suggest that cystine/cysteine directly fuels the thioredoxin system to prevent BSO-induced ferroptosis^13^, our data indicate that cysteine incorporation in CoA biosynthesis is essential for regulating mitochondrial thioredoxin system to protect against ferroptosis.

### Protein CoAlation on TXNRD2 determines its enzymatic activity

While the protein levels of thioredoxin reductase remain unchanged, the enzymatic activity of thioredoxin reductase was significantly inhibited by erastin, and such inhibition could be rescued by CoA, suggesting involvement of post-translational modifications (Fig. 2B, Supplemental Fig. 2O). Recently, CoA has been demonstrated to modify proteins by covalent attachment (CoAlation) to the thiol groups of cysteine residues of various cellular proteins, thereby regulating their enzymatic activities^6^. Interestingly, TXNRD2 is one of the potential CoAlated hits^6^. Thus, we raised a monoclonal antibody that specifically recognized CoA and CoAlation proteins (Supplementary Fig. 3A). By supplementing CoA in HT-1080 cells overexpressing TXNRD2-V5 and TXN2-myc, we found that CoA increased the interaction between TXNRD2 and TXN2. Importantly, TXNRD2 pulldown showed increased CoAlation upon CoA supplementation (Fig. 3A). To verify whether TXNRD2 can be CoAlated *in vitro*, we purified V5 tagged TXNRD2 protein from HT-1080 cells overexpressing TXNRD2-V5 and incubated with oxidized CoA using a previously published protocol^6^. The purified proteins were then separated using native gel and blotted with anti-CoA antibody. We confirmed that TXNRD2 was indeed robustly CoAlated, and treatment with oxidized CoA led to a significant shift in migration behaviors, suggesting structural changes (Supplemental Fig. 3B). Next, we ran oxidized CoA-treated TXNRD2 in non-reducing gel and stained with anti-CoA antibody. Consistently, we found that treatment with oxidized CoA dramatically increased TXNRD2 CoAlation. This effect was abolished by dithiothreitol (DTT), which disrupted disulfide bond of CoAlation (Fig. 3B). Next, we examined whether CoAlation of TXNRD2 affects its enzymatic activity (Fig. 3C). Indeed, using the *in vitro* thioredoxin reductase assay, we found that CoAlation on TXNRD2 protein increased enzymatic activity, and this effect could be completely inhibited by the thioredoxin reductase inhibitor: auranofin^27^(Fig. 3C).

**Fig. 3.**
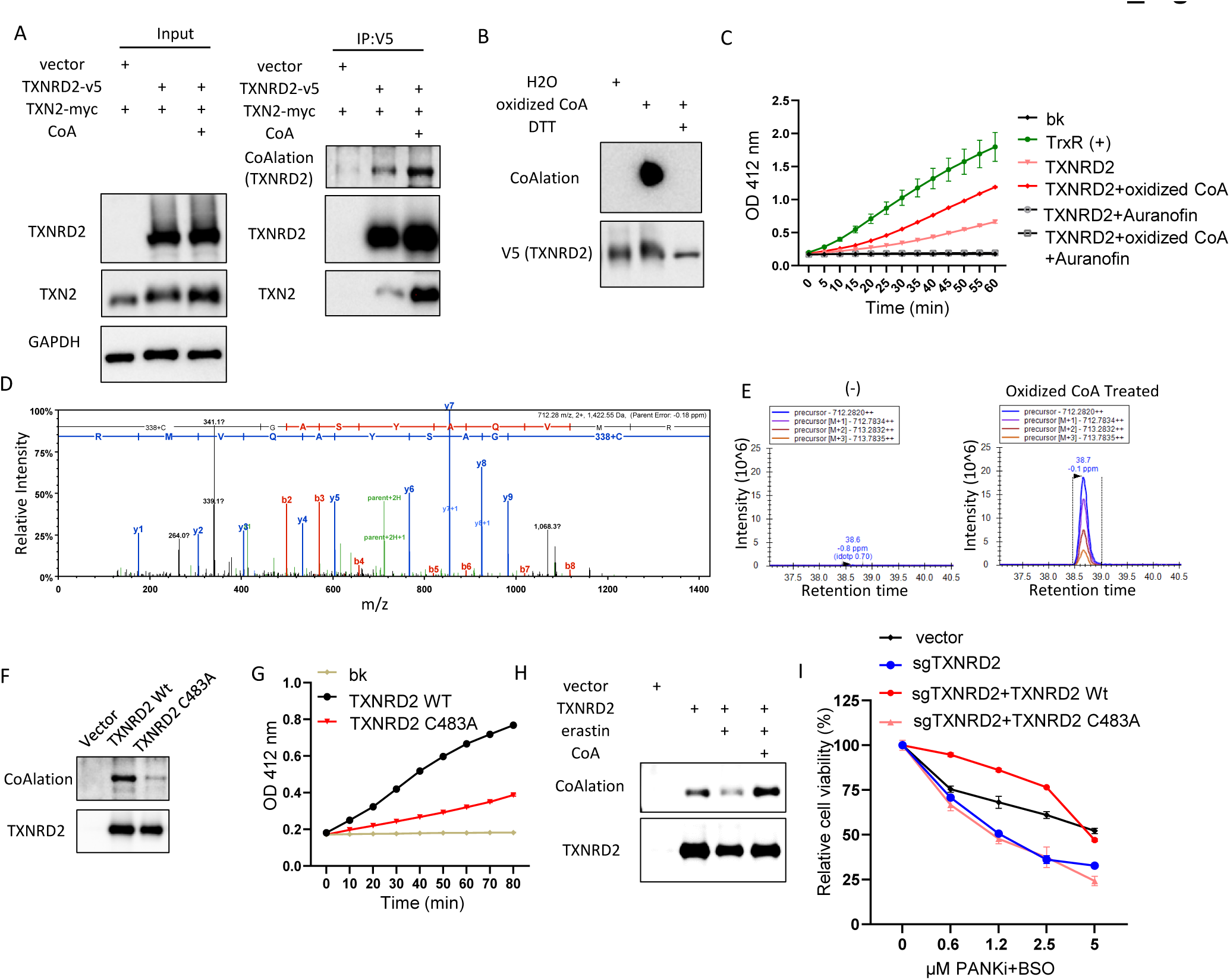
CoAlation of Cys-483 on TXNRD2 protein regulates the thioredoxin reductase activity. (**A**) CoA supplement increased the CoAlation of TXNRD2 and its interaction with TXN2. HT-1080 cells overexpressing TXNRD2 and TXN2 were supplemented with CoA (100 μM, 18 hours), and the TXNRD2-TXN2 interaction was evaluated by co-immunoprecipitation. (**B**) The specificity of the CoAlation antibody was verified by DTT, which abolishs the disulfide bond. V5-tag purified TXNRD2 was treated with oxidized CoA or combined with DTT, resolved on non-reducing PAGE, and blotted for Western blots. (**C**) CoAlation increased thioredoxin reductase activity of TXNRD2. The enzymatic activities of thioredoxin reductase of purified TXNRD2 protein with or without CoAlation or in combination with TXNRD inhibitor (Auranofin). The graph also included the background (bk) and positive control of thioredoxin reductase (TrxR(+)). (**D**) Tandem mass spectrum of CoA-modified peptide “C*GASYAQVMR” within TXNRD2. (**E**) Extracted Ion chromatograms (EIC) of the 2+ precursor ion corresponding to “C*GASYAQVMR in the DMSO control non-CoA sample and CoA treated sample demonstrating the lack of modified peptide in the DMSO control. (**F**) Replacing Cys-483 with Alanine abolished most of the CoAlation on the TXNRD2 protein. Purified wild-type or C483A mutant TXNRD2 proteins were resolved on non-reducing PAGE and Western blots for protein CoAlation. (**G**) Replacing Cys-483 with Alanine in TXNRD2 (C483A) protein abolished thioredoxin reductase activity. bk, background. (**H**) Erastin reduced the CoAlation on TXNRD2, which was restored by CoA treatment. HT-1080 cells overexpressing TXNRD2 were treated with erastin (2 μM) or in combination with CoA (100 μM) for 16 hours. TXNRD2 proteins were purified by V5 tag and Western blots for CoAlation. (**I**) Cys-483 on TXNRD2 determines its function against ferroptosis induced by the combination of PANKi and BSO. HT-1080 cells were transduced with lentiviral sgRNA against TXNRD2 to knock out endogenous TXNRD2 expression and overexpressed with TXNRD2 wild-type or C483A mutant with synonymous mutation to avoid targeting by TXNRD2 sgRNA. These cell lines were treated with the combination of BSO (300 μM) and various concentrations of PANKi for Cell-Titer Glo assay.

To identify the particular CoAlated residue(s) on TXNRD2 protein, we performed a bottom-up mass spectrometry experiment on oxidized CoA-treated tryptic digested TXNRD2 protein (Fig. 3D-E, Supplementary Fig. 3C). Compared to untreated control, we identified a dynamic mass modification of +338.07 Da corresponding to fragmented CoA on Cys-483 of TXNRD2 (Fig. 3D-E, Supplementary Fig. 3C). Although the searches also included full-length CoA (+762.08 Da), full-length CoA was not identified in these data, possibly due to the instability of the intact structure in the gas phase causing in-source fragmentation between phosphate groups. The 3D structure of TXNRD2 has been solved^28^, and Cys-483 is highly conserved during evolution and does not participate in the formation of intramolecular disulfide bonds^28^. Cys-483 has also been postulated to have a potential redox regulatory role^28^. To identify the function of this CoAlation residue, we used site-directed mutagenesis to replace Cys-483 with alanine (C483A) on TXNRD2 and transduced wild-type and the C483A mutant into HT-1080 cells (Fig. 3F). Indeed, after protein purification, we found that wild-type TXNRD2 protein had robust endogenous levels of CoAlation signal, which was mostly abolished in C483A mutant (Fig. 3F). Also, this endogenous CoAlation signal can be removed by beta-mercaptoethanol (beta-ME) treatment, thereby disassociating CoA from modified Cys-483 and further demonstrating the specificity on this CoAlation signal (Supplementary Fig. 3D). Therefore, we tested whether CoAlation on Cys-483 determines the enzymatic activity of TXNRD2 (Fig. 3G). Compared with the wild-type TXNRD2 protein, the thioredoxin reductase activity of the C483A CoAlation-null mutant was significantly decreased (Fig. 3G). Collectively, these data suggest that CoAlation on Cys-483 of TXNRD2 protein determines its thioredoxin reductase activity.

Next, we tested whether CoAlation on TXNRD2 protein could be modulated by cystine/cysteine availability. In TXNRD2-overexpressed HT-1080 cells treated with erastin alone or in combination with CoA, we found that erastin reduced CoAlation on purified TXNRD2, which could be rescued by CoA supplementation (Fig. 3H). Similarly, both inhibition of CoA biosynthesis by PANKi or direct inhibition of TXNRD2 by auranofin reduced the CoAlation of TXNRD2 protein (Supplementary Fig. 3E-F). To demonstrate that CoAlation on TXNRD2 determines cell viability, we performed CRISPR to knock out endogenous TXNRD2. By treating these cells with a combination of BSO and PANKi, we found that knockdown of TXNRD2 sensitized cells to ferroptosis. Next, the CRISPR-resistant wild-type or C483A mutant TXNRD2 was re-expressed in the TXNRD2-null cells (Supplementary Fig. 3G). The BSO/PANKi sensitizing effects in the TXNRD2-null cells were fully rescued by wild-type TXNRD2 but not C483A mutant (Fig. 3I). These data suggest that CoAlation on the Cys-483 of TXNRD2 protein regulates its enzymatic activity and ferroptosis sensitivity.

### Disruption of CoA metabolism by PANK inhibition leads to mitochondrial lipid peroxidation and ferroptosis

Mutations in human genes *PANK2* or *COASY*, two genes that encode proteins involved in the CoA biosynthesis pathway, lead to NBIA^2^. In particular, mutations in the *PANK2* lead to PKAN^2^. The mammalian genome encodes four isozymes possessing pantothenate kinase activity (PANK1α, PANK1β, PANK2, and PANK3). Their tissue expression patterns, and subcellular localization have been extensively documented^3^. However, the respective cellular roles of each PANK protein and the unique cellular role of *PANK2* mutations in tissue-specific vulnerability leading to PKAN are still unclear. By selectively targeting the glutathione synthesis using BSO, we found that suppressing the mitochondrial thioredoxin system by inhibiting PANK or knocking down COASY significantly increased ferroptosis (Fig. 2I-J) and mitochondrial lipid peroxidation (Fig. 2K-M). To investigate the role of individual PANKs in CoA-directed thioredoxin regulation, we knocked down individual PANK proteins in HT-1080 cells and found that only PANK1 knockdown increased BSO-induced ferroptosis (Fig. 4A, Supplementary Fig. 4A). Surprisingly, by using BSO to selectively target glutathione system, while the fibroblasts from healthy individuals were unaffected, the fibroblasts from PKAN patients showed a significant decrease in cell viability (Supplementary Fig. 4B). Importantly, CoA supplementation was able to rescue the BSO-induced cell death in fibroblasts from PKAN patients (Supplementary Fig. 4C). Next, in HT-1080 cells, the knockdown of PANK1 and PANK2, but not PANK3, increased mitochondrial lipid peroxidation to the same level as COASY knockdown (Fig. 4B). Furthermore, we compared mitochondrial ROS and mitochondrial lipid peroxidation of fibroblasts between PKAN patients and healthy individuals (Fig. 4C-D, Supplementary Fig. 4D-E). Surprisingly, we observed increased mitochondrial lipid ROS and mitochondrial lipid peroxidation in fibroblasts from PKAN patients (Fig. 4C-D, Supplementary Fig. 4D-E). While *PANK2* mutation/knockdown may exhibit different responses in cell viability to the stress of BSO-mediated GSH depletion, these data suggest a consistent increase in mitochondrial lipid peroxidation upon *PANK2* mutation/knockdown in PKAN fibroblast and HT-1080 cells.

**Fig. 4.**
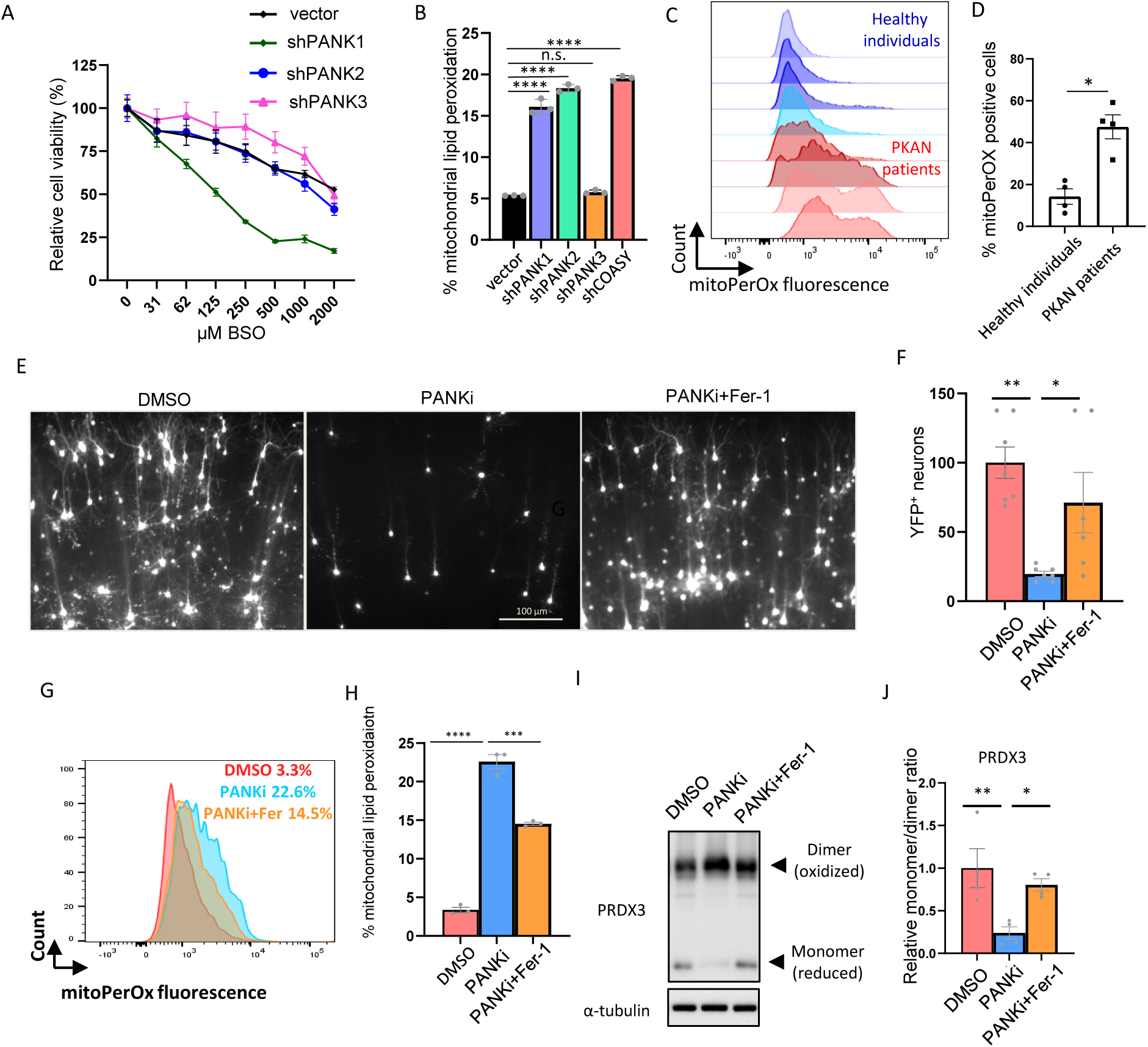
Disruption of CoA biosynthesis in PKAN fibroblasts and OBSC leads to mitochondrial lipid peroxidation. (**A**) *PANK1* knockdown sensitized HT-1080 cells to BSO treatment. HT-1080 cells with shRNA targeting *PANK1*, *PANK2*, or *PANK3* were treated with different doses of BSO for three days for Cell-Titer Glo assay. (**B**) *PANK1*, *PANK2*, and *COASY* knockdown by shRNA in HT-1080 cells showed an increase in mitochondrial lipid peroxidation. (**C-D**) PKAN, when compared with healthy, fibroblasts showed elevated mitochondrial lipid peroxidation. Four pairs of fibroblasts from PKAN patients and unaffected individuals were stained with the sensor of mitochondrial lipid peroxidation (mitoPerOx) (**C**) and the quantification of % mitochondrial lipid peroxidation positive cells (**D**). (**E-F**) The cell death triggered by inhibiting CoA using PANKi in OBSC was rescued by ferrostatin-1. OBSC transfected with YFP were treated with PANKi (2.5 μM) or in combination with ferrostatin-1 (Fer-1, 2 μM) for one day, and the neuron numbers were accessed by fluorescence microscopy (**E**) and quantified in (**F**). (**G-H**) The elevated mitochondrial lipid peroxidation by PANKi was rescued by ferrostatin-1. After 1 day of treatment with PANKi (2.5 μM) and Fer-1 (2 μM), OBSC was disassociated and stained with mitoPerOx (**G**) for quantification (**H**). (**I-J**) PANKi treatment of brain slices repressed the levels of reduced and active form of PRDX3, which was rescued by Fer-1. OBSC treated with PANKi (2.5 μM) or in combination with Fer-1(2 μM) were homogenized to blot for mitochondrial PRDX3 (**I**) and quantification (**J**).

To bridge the gap between cell-based assays and whole animal models, organotypic brain slice culture (OBSC) has been developed to investigate the mechanisms of diseases and accelerate drug development^29–31^. OBSC also serves as a model system to study ferroptosis effects in neurons^32^. To mimic the human CoA metabolic defect, we treated rat OBSC with PANKi to inhibit CoA biosynthesis. By monitoring the number and viability of neurons by transfecting OBSC with YFP, we observed that the number of neurons was reduced after PANKi treatment, which could be rescued by the ferroptosis inhibitor, ferrostatin-1 (Fig. 4E-F). These data suggest that inhibition of the thioredoxin system by PANKi itself is sufficient to trigger ferroptosis in the OBSC model. Subsequently, we measured mitochondrial lipid peroxidation and found that mitochondrial lipid peroxidation was consistently increased upon PANKi treatment, which could be abolished through the concurrent application of ferroptosis inhibitor: ferrostatin-1 (Fig. 4G-H). These data suggest that the PANKi triggered mitochondrial lipid peroxidation in neurons in the OBSC model. Finally, we measured the redox status of the mitochondrial form of peroxiredoxin (PRDX3) and found that PANKi downregulated the levels of reduced monomers, which were restored by ferrostain-1 treatment (Fig. 4I-J). While PANKi treatment itself did not induce ferroptosis in the six cancer cell lines (Supplementary Fig. 4F), our data suggest that inhibiting CoA biosynthesis may trigger ferroptosis by inducing mitochondrial lipid peroxidation in rat neurons in the OBSC model.

## Discussion

CoA was first identified as a ferroptosis inhibitor in p53-mutant cells with increased CoA levels that protect against erastin and glutamate-induced ferroptosis^14^. In pancreatic tumor cells, CoA was proposed to protect ferroptosis by enhancing the production of coenzyme Q_10_, ^8^. In this study, we found that CoA serves as a protective agent against ferroptosis induced by class I, but not other classes of FINs in multiple cancer cell lines (Fig. 1C, Supplementary Fig. 1A-E). Previous studies have found that class I FIN requires mitochondria and their lipid peroxidation to trigger ferroptosis via an unknown mechanism^10,11^. Curiously, while BSO significantly reduces glutathione synthesis, BSO is a considerably less potent FIN to trigger ferroptosis. One group proposed that GSH inhibition by BSO led to the accumulation of intracellular cysteine levels, which were used by the thioredoxin system to suppress ferroptosis^13^. However, the mechanism by which cysteine regulates the thioredoxin system has not been fully elucidated^13^. Instead of direct fueling of thioredoxin system by cysteine, our findings suggest that cysteine is incorporated for the biosynthesis of CoA, which protects cells from ferroptosis by CoAlation of TXNRD2 to maintain a mitochondrial thioredoxin function and prevent mitochondrial lipid peroxidation. Thus, xCT-cysteine-CoA-TXNRD2 is an important independent branch of the ferroptosis defense mechanism, distinct from the xCT-GSH-GPX4 axis, that neutralizes mitochondrial lipid peroxidation.

Protein CoAlation was first identified in 2017 as a reversible post-translational modification in eukaryotic and prokaryotic cells in response to oxidizing environments and metabolic stress^6,33^. So far, CoAlation has been found to inhibit the activity of modified proteins implicated in metabolic or signaling pathways^7^. The inhibitory effects of CoAlation have been demonstrated in aconitase, creatine kinase, pyruvate dehydrogenase kinase 2, glyceraldehyde 3-phosphate dehydrogenase, hydroxymethylglutaryl-CoA synthase, Aurora A kinase, and metastasis suppressor protein NME1^6,7,34–36^. In contrast to these reported inhibitory effects, we found that CoAlation on Cys483 of TXNRD2 increased its thioredoxin reductase activity, which may involve an allosteric mode of activation (Fig. 3C, G). Importantly, CoAlation levels on TXNRD2 respond to the availability of cysteine and CoA as a nutrient-sensing mechanism (Fig.3H, Supplementary Fig. 3E). Furthermore, direct inhibition of TXNRD2 with auranofin reduced TXNRD2 CoAlation and abolished CoA protection against ferroptosis. (Fig. 2A, Supplementary Fig 3F). Upon the dual inhibition of glutathione and CoA synthesis (BSO and PANKi), CoAlation on TXNRD2 determined cell viability (Fig. 3I). Importantly, Cys483 is highly conserved among mammalian TXNRDs^28^ and is surface-exposed. Therefore, this residue has been postulated to have a potential redox regulatory role^28^. Our findings confirm this prediction that CoAlation on Cys483 is a nutrient-sensing mechanism that adjusts the enzymatic activity of TXNRD2 and ferroptosis sensitivity based on the levels of CoA. Given that Cys483 is highly conserved during evolution ^28^, redox status regulation by CoAlation may have broader impacts on other TXNRDs across different species as CoA biosynthesis is an ancient pathway conserved across different domains of life^37^.

NBIA is a group of rare neurodegenerative disorders with increased basal ganglia iron on brain magnetic resonance imaging^38^. NBIA is associated with mutations in different enzymes involved in the CoA biosynthesis, including *PANK2* and *COASY*. Mutations in *PANK2*, a key mitochondrial enzyme involved in CoA biosynthesis, account for approximately half of NBIA cases^38^. Given that iron is essential for ferroptosis via the Fenton reaction, it is reasonable to speculate on a relationship between ferroptosis and PKAN/NBIA. Consistently, PKAN astrocytes were found to be prone to ferroptosis with higher oxidized proteins and malondialdehyde^39^. However, a direct connection between disrupted CoA metabolism and ferroptosis during PKAN is still lacking. In this study, we found that *PANK2* knockdown increased mitochondrial lipid peroxidation in HT-1080 cells (Fig. 4B). Even though PKAN patients showed no reduction in CoA levels in fibroblasts^3^, we still observed increased mitochondrial lipid peroxidation (Fig. 4C-D) that may predispose these cells to ferroptosis. Indeed, fibroblasts from PKAN patients are more sensitive to BSO-mediated glutathione depletion (Supplementary Fig 4B). We also found that OBSC treated with PANKi exhibited many features associated with PKAN/NBIA, including a reduced number of neurons, increased mitochondrial lipid peroxidation, and disrupted thioredoxin systems (Fig. 4E-J). Importantly, the application of ferrostatin-1 to rescue these phenotypes implies that PANKi-induced cell death is caused by ferroptosis (Fig. 4E-J). These data also suggest that we have developed an OBSC model targeting all PANKs to complement current models of PKAN and NBIA. With the development of PANK2-specific inhibitor in the future or genetic removal of *PANK2*, this OBSC model can be further developed to reveal the mechanistic underpinnings and therapeutic agents. Inhibiting ferroptosis may have significant therapeutic implications.

## Methods

### Mass spectrometry

CoA and acetyl-CoA were extracted from 30 million cells per sample using previously published methods^40^. The cell pellets were treated with 300 µL of ice-cold 0.1M KH2PO4. Methanol containing 10% acetic acid was added to the cell pellets, followed by solid phase extraction (SPE). For the SPE process, 1 ml ion exchange cartridge packed with 100 mg of 2– 2(pyridyl)ethyl silica gel (Sigma) was activated with 1 ml of methanol and equilibrated with 1 ml of buffer A (methanol/H_2_O 1:1, containing 5% acetic acid). The sample extract was loaded onto the cartridge, and additional steps with buffer A, buffer B (methanol/H_2_O 1:1, 50 mM ammonium formate), buffer C (methanol/H_2_O 3:1, 50 mM ammonium formate), and methanol were performed. The eluent (3 ml) was collected and dried using N_2_ gas. The resulting dried residue was re-suspended in 100 µl of H_2_O for LC-MS/MS analysis^40,41^.

For identification of CoAlation site on TXNRD2, HT-1080 cells with TXNRD2-v5 over-expression was lysed with NP-40 buffer with 25 mM N-ethylmaleimide (NEM, 23030, Thermo Fisher) and protease inhibitor (04693116001, Roche) in 4 degree with constant shaking. The lysates were then centrifuged at 21,000 g for 10min, and the supernatant was purified by V5-tagged Protein Purification Kit (3317, MBL) and eluted by v5 peptide. Purified TXNRD2 was incubated with 10 mM oxidized CoA in 50 mM Tris-HCl (pH 7.5) at room temperature for 30 min. The samples with/without oxidized CoA treatment were then resolved on a 12% non-reducing SDS PAGE. After colloidal blue staining (LC6025, Thermo fisher), the bands with ∼50kD were cut for Mass spectrometry.

The bands obtained from SDS-PAGE gels underwent in-gel tryptic digestion following established protocols. Post-digestion, peptides were dehydrated and then reconstituted in a solution containing 0.2% formic acid and 2% acetonitrile at a volume of 12 uL. Each resulting sample underwent chromatographic separation using a Waters MClass UPLC, utilizing a 1.7 µm HSS T3 C18 75 µm I.D. X 250 mm reversed-phase column (NanoFlow data). The mobile phase for chromatography consisted of two components: (A) 0.1% formic acid in water and (B) 0.1% formic acid in acetonitrile. Injection of 3 µL of the sample commenced the process, with peptides initially captured for 3 minutes on a 5 µm Symmetry C18 180 µm I.D. X 20 mm column at a rate of 5 µl/min, with a composition of 99.9% A. Following this, an analytical column was engaged, and a linear elution gradient ranging from 5% B to 40% B occurred over 60 minutes at a flow rate of 400 nL/min. The Fusion Lumos mass spectrometer (Thermo) was connected to the analytical column via an electrospray interface, functioning in a data-dependent acquisition mode. The instrument settings included a precursor MS scan from m/z 375-1500 at R=120,000 (target AGC 2e5, max IT 50 ms), with subsequent MS/MS spectra acquired in the ion trap (target AGC 1e4, max IT 100 ms). HCD energy settings were consistently maintained at 30v for all experiments, and a dynamic exclusion of 20 seconds was implemented to avoid re-analysis of previously fragmented precursor ions.

The LC-MS/MS data files obtained were processed using Proteome Discoverer 3.0 (Thermo Scientific) and subsequently subjected to independent Sequest database searches against a comprehensive Human protein database. This database included both forward (20260 entries) and reverse entries for each protein. The search parameters allowed for 2 ppm tolerance for precursor ions and 0.8 Da tolerance for product ions, employing trypsin specificity with allowance for up to two missed cleavages. Dynamic mass modifications were defined for Cleaved CoA (+338.07 Da on C) and full-length CoA (+762.08 Da on C). All spectra derived from the searches were imported into Scaffold (v5.3, Proteome Software), where scoring thresholds were adjusted to maintain a peptide false discovery rate of 1%, utilizing the PeptideProphet algorithm. Additionally, the raw data was imported into Skyline (MacCoss Lab, UWash) for the purpose of measuring extracted ion chromatograms of modified peptides.

### Chemicals

Auranofin (A6733, Sigma); ferroptocide (F1293, TCI); PANKi (31002, Cayman); CoA (F15115, Astatech); erastin (5499, Bio-techne); BSO (B2515, Sigma); RSL3 (19288, Cayman); ML162 (20455, Cayman); FIN56 (25180, Cayman); ferrostatin-1 (17729, Cayman); liproxstatin-1 (17730, Cayman); Trolox (218940010, Thermo Fisher); lovastatin (S2061, Selleckchem); simvastatin (S1792, Selleckchem); iFSP1 (29483, Cayman); brequinar (S3565, Selleckchem); Teriflunomide (S4169, Selleckchem); Methotrexate (S1210, Selleckchem); Compound C (S7306, Selleckchem); TOFA (S6690, Selleckchem); Etomoxir sodium salt (S8244, Selleckchem); 3-Deazaadenosine (HY-W013332A, Medchemexpress); 3-Deazaneplanocin A (HY-12186, Medchemexpress); Adenosine dialdehyde (HY-123055, Medchemexpress).

### In vitro assay

To determine thioredoxin activity in cell lysate, after the treatment of erastin, CoA, or in combination, 30 million cells per sample were washed with PBS and homogenized in 500μl of cold buffer (100 mM Tris-HCl, pH7.5, 1 mM EDTA) with protease inhibitor. After 15 minutes of centrifugation (10,000g, 4 degree Celsius), the thioredoxin activity in the supernatant was determined by Thioredoxin Fluorometric Activity Assay Kit (500228, Cayman). To determine thioredoxin reductase activity in cell lysate, after the treatment of erastin, CoA, or in combination, 30 million cells per sample were washed with PBS and homogenized in 500μl of cold buffer (50 mM potassium phosphate, pH 7.4, 1 mM EDTA) with protease inhibitor. After 15 minutes of centrifugation (10,000g, 4 degree Celsius), the thioredoxin activity in the supernatant was determined by Thioredoxin Reductase Colorimetric Assay Kit (10007892, Cayman). To determine thioredoxin reductase activity in purified TXNRD2 protein, HT-1080 cells transduced with TXNRD2 Wild-type or C483A mutant were lysed with NP-40 buffer with protease inhibitor and 25 mM NEM. After centrifugation, the supernatant was purified by V5-tagged Purification Kit Ver.2 (3317, MBL). The purified TXNRD2 protein was resolved by non-reducing SDS PAGE to confirm equal amount of input for Thioredoxin Reductase Colorimetric Assay Kit (10007892, Cayman).

### Cell culture

HT-1080, HEK-293, 786-O, H1975, RCC4, and MDA-MB-231 cells were sourced from the Cell Culture Facility at Duke University (Durham, NC, USA). Prior to freezing, these cell lines were authenticated using STR DNA profiling to ensure their identity and confirmed to be free from mycoplasma contamination by the Cell Culture Facility. The cells were cultured for a duration of less than 6 months. Human fibroblasts from healthy individuals were obtained from Coriell Institute for Medical Research. The fibroblasts from PKAN patients were gifts from Dr. Susan J. Hayflick. All fibroblast collection are part of Duke’s IRB-approved protocol Pro00101047. All cell lines were maintained in a humidified incubator at 37°C with 5% CO2 in DMEM (GIBCO-11995) supplemented with 10% heat-inactivated fetal bovine serum (#10082147, ThermoFisher) and antibiotics (streptomycin, 10,000 UI/ml and penicillin, 10,000 UI/ml, #15140122, ThermoFisher).

### Constructs and lentivirus viral infections

Small interfering RNAs (siRNAs) targeting human *COASY*, *TXN1*, *TXN2*, *ACSL3*, and *ACSL4* RNA were obtained from Dharmacon (D-006751-01, D-006751-02, M-006340-01-0005, M-017448-00-0005, M-010061-00-0005, M-009364-00-0005). The guide RNA targeting TXNRD2 was purchased from Sigma (HSPD0000063630 in GeCKOv2 all-in-one lentiviral plasmid). The TXNRD2 cDNA in PLX304 was procured from DNASU (HsCD00446697). To generate guide RNA-resistant TXNRD2 (with synonymous mutation) and point mutant (TXNRD2 C483A), we employed the QuikChange II XL Site-Directed Mutagenesis Kit (#200521, Agilent). Lentivirus expressing specific constructs was produced by transfecting HEK-293 cells in 6-well plates with a 1:1:0.1 ratio of lentiviral vector:pMD2.G:psPAX2 using the TransIT-LT1 transfection reagent (Mirus). Subsequently, the lentivirus was filtered through a cellulose acetate membrane (0.45 µm, #28145-481, VWR), and 250 µl of media containing the lentivirus were added to a 60mm dish of the indicated cells along with polybrene (8 µg/ml) and selected with puromycin or blasticidin.

### Cell viability and cytotoxicity

Cell viability was assessed using the CellTiter-Glo luminescent cell viability assay (Promega) according to the manufacturer’s instructions. Briefly, 15 µl of CellTiter-Glo substrate was added to cells cultured in a 96-well plate with 100 µl of media, followed by 10 minutes of shaking. The signal intensity was measured using a chemiluminescence plate reader. For quantification of cell death, the CellTox Green assay (Promega) was performed. The dye was added to the media at a 1:1000 dilution, and cell death was quantified using a fluorescence plate reader. To determine the level of GSH (glutathione), the GSH/GSSG-Glo Assay (Promega) was conducted according to the manufacturer’s guidelines. Briefly, the samples in a 96-well plate were treated with a lysis reagent specific to each measurement set: total glutathione lysis reagent for total glutathione measurement and oxidized glutathione lysis reagent for GSSG measurement. Afterward, Luciferin Generation Reagent was added to all wells, and the assays were mixed and incubated for 30 minutes. Following this, Luciferin Detection Reagent was added to all wells, and luminescence was measured after a 15-minute incubation. GSH/GSSG ratios were directly calculated from the luminescence measurements (in relative light units, RLU).

### Western blots

Protein concentrations were quantified by BCA assay (#23227, ThermoFisher). After protein extraction, the samples were loaded on 12% native, non-reducing, or reducing SDS-PAGE gels, transferred to PDVF membrane, blocked with 5% non-fat milk in 1xTBST, incubated with primary antibodies overnight at 4°C. Primary antibodies: COASY (1:1000, sc-393812, Santa Cruz); PRDX1 (1:1000, 15816-1-AP, Proteintech); PRDX3 (1:1000, 10664-1-AP, Thermo Fisher); GAPDH (1:2000, sc-25778, Santa Cruz); TXNRD1 (1:1000, 11117-1-AP, Santa Cruz); TXNRD2 (1:1000, PA529458, Thermo Fisher); v5 (1:1000, MA5-15253, ThermoFisher); alpha Tubulin (1:1000, sc-32293, Santa Cruz). CoAlation antibody is a gift from Dr. Chen-Yong Lin (PT-mAb-CoA, ProtTech). TXNRD2 protein was purified from HT-1080 cells overexpressing TXNRD2-v5 by V5-tagged Protein Purification Kit Ver.2 (3317, MBL). For TXNRD2 purification, HT-1080 cells were lysed by NP-40 buffer with 25 mM N-ethylmaleimide (NEM, 23030, Thermo Fisher) and protease inhibitor (04693116001, Roche) in 4 degree with constant shaking. The lysates were then centrifuged at 21,000 g for 10min, and the supernatant was purified by V5-tagged Protein Purification Kit. For PRDX blots, the cells were washed with PBS and incubated in NEM buffer (HEPES, pH 7.4 40mM, NaCl 50 mM, EDTA 1mM, EGTA 1mM, NEM 100 mM, protease inhibitor) for 10 min. CHAPS (1%) was then added to lyse the cells for quantification.

### Quantitative real-time PCR

The RNA extraction and purification were carried out using the RNeasy Mini Kit (Qiagen) following the manufacturer’s recommended protocol. Subsequently, reverse transcription to cDNA was performed using random hexamers and SuperScript IV reverse transcriptase (Invitrogen). For quantitative real-time PCR analysis, the cDNA was combined with primers and Power SYBR Green PCR Mix (Applied Biosystems), and the reactions were run on a StepOnePlus Real-time PCR system (Applied Biosystems). Each sample was technically triplicated to obtain the mean +/- standard error of the mean (SEM). The data presented are representative of a minimum of two independent experimental repetitions. Primer sequences are listed in Supplemental Table 1.

### Lipid peroxidation assay

Lipid peroxidation was assessed using C11-BODIPY staining following the manufacturer’s instructions (D3861, ThermoFisher Scientific). Briefly, cells were exposed to either a vehicle control or treatments for 16 hours. Subsequently, the medium was replaced with a 10µM C11-BODIPY-containing medium and incubated for 1 hour. After harvesting, washing, and resuspension in PBS containing 1% BSA, the levels of lipid peroxidation were quantified using flow cytometry (FACSCanto TM II, BD Biosciences). Similarly, for mitochondrial lipid peroxidation, mitoPerOx^43^ (200 nM,18798, Cayman) containing medium was incubated with cells for 30 min.

### Organotypic brain slice cultures

The experiment involved the use of CD Sprague-Dawley rat pups at postnatal day 10, obtained from Charles River (Wilmington, MA). The rat brains were sliced into 250 μm coronal sections using Vibratomes from Vibratome Co. (St. Louis, MO) in chilled medium baths. Each rat brain yielded approximately 6 slices, which were further divided into “hemi-coronal” slices. These slices were individually placed in multi-well plates, resulting in 12 brain slice assays per rat brain.

To establish the brain slice assays, the slices were explanted in an interface configuration using culture medium containing 15% heat-inactivated horse serum, 10 mM KCl, 10 mM HEPES, 100 U/ml penicillin/streptomycin, 1 mM MEM sodium pyruvate, and 1 mM L-glutamine in Neurobasal A (Invitrogen, Carlsbad, CA). Before use, the medium was filter sterilized at 0.22 μm.

The 12-well plates used for plating the slices provided culture support through a low concentration of agarose (0.5%; J.T. Baker, Phillipsburg, NJ). The slice cultures were maintained in humidified incubators at 32 °C under 5% CO2.

To label healthy neurons, YFP was transfected using a biolistic device (Bio-Rad Helios Gene Gun). Following the treatment of PANKi or a combination of PANKi with ferrostatin-1, neurons expressing YFP were observed and visualized under fluorescence microscopy after 1 day of incubation.

All work performed on Sprague-Dawley pup for OBSC was approved by the Institutional Animal Care and Use Committee at Duke University.

### Statistical analysis

The bar graphs display individual data points, each representing the number of biological replicates as indicated. The line graph includes the number of biological replicates specified in the figure legends. All data are presented as the mean +/- the standard error of the mean (SEM). Statistical significance was determined using Graphpad software with one-way ANOVA and Tukey’s multiple comparisons, two-way ANOVA with Dunnett’s multiple comparisons, or a two-tailed Student’s t-test, as appropriate. The error bars on the graphs represent the SEM, and significance between samples is denoted as *p < 0.05, **p < 0.01, ***p < 0.001, and ****p < 0.0001.

### Data availability

The authors are prepared to provide all data and reagents supporting the study’s findings upon a reasonable request.

## Author contributions

C.C.L. and J.T.C. conceived the experiments and wrote the manuscript. C.C.L. performed the majority of the experiments. J.T.C. supervised the work. Y.T.L., S.Y.C., Y.S., Y.C., D.E.D., E.J.S., S.F., S.Y.J., V.F., and G.Z. collaborated in the discussion and experiments. S.J.H. and I.G. provided critical feedback.

## Acknowledgments

We are grateful for technical support from the members of the Chi lab and Zih-Syuan Wu from National Defense Medical Center. We acknowledge the financial support in part by DCI Pilot Project, DOD grants (W81XWH-17-1-0143, W81XWH-15-1-0486, W81XWH-19-1-0842, W81XWH-20-1-0907) and NIH grants (R01GM124062, 1R01NS111588-01A1, 1R21-AI149205).

---

**Supplementary Fig. 1.**
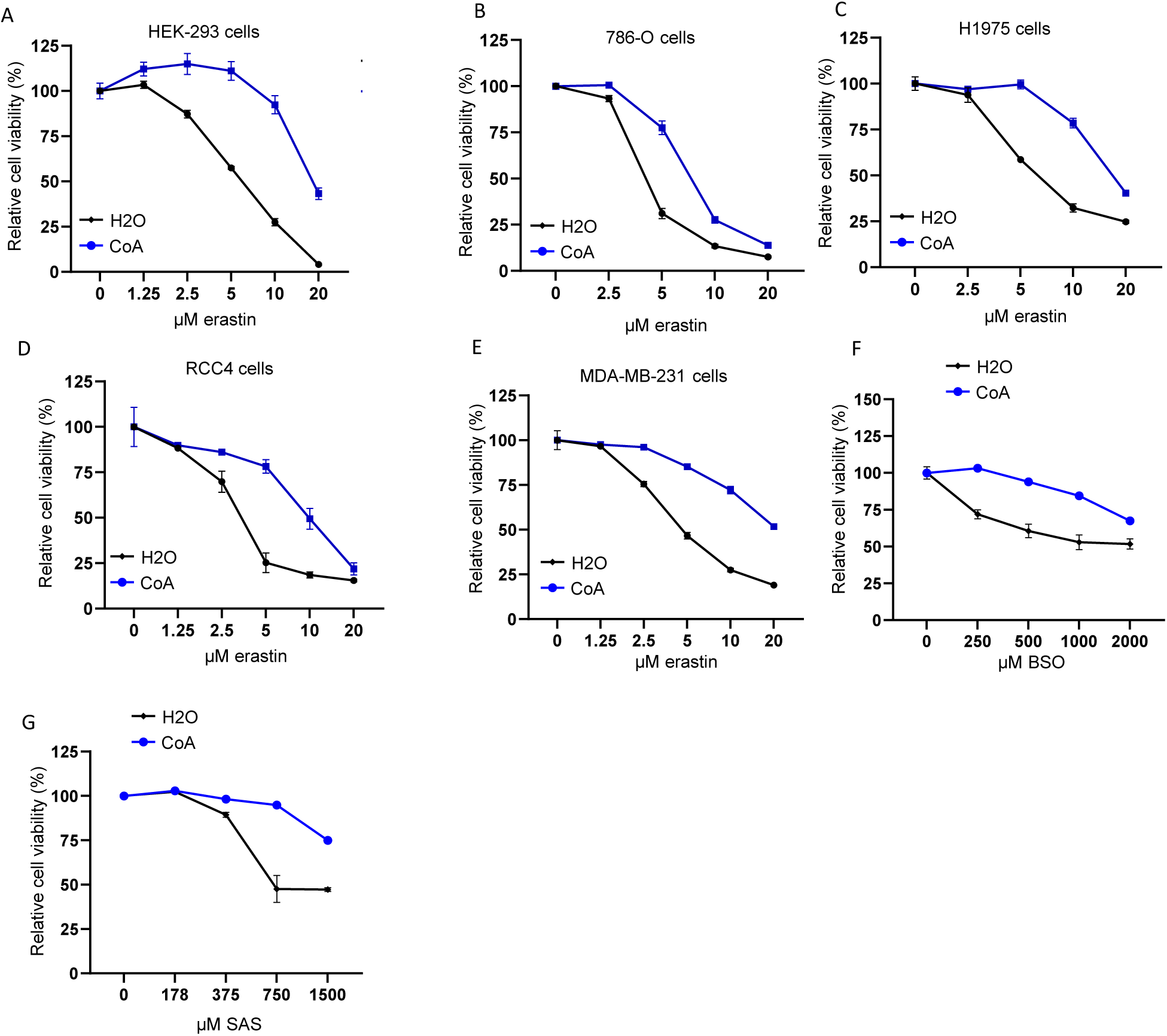
The effects of CoA on the ferroptosis of various cancer cell lines induced by different FINs. (**A-E**) HEK-293 (**A**), 786-O (**B**), H1975 (**C**), RCC4 (**D**), and MDA-MB-231 cells (**E**) were treated with CoA (100 μM) and in combination with varying doses of erastin for 18 hours. The cell viability was quantified by Cell-Titer Glo assay. (**F-G**) CoA (100 μM) inhibited class 1 FIN-induced ferroptosis in HT-1080 cells including BSO (3 days) (**F**) and sulfasalazine (SAS, 20 hours) (**G**).

**Supplementary Fig. 2.**
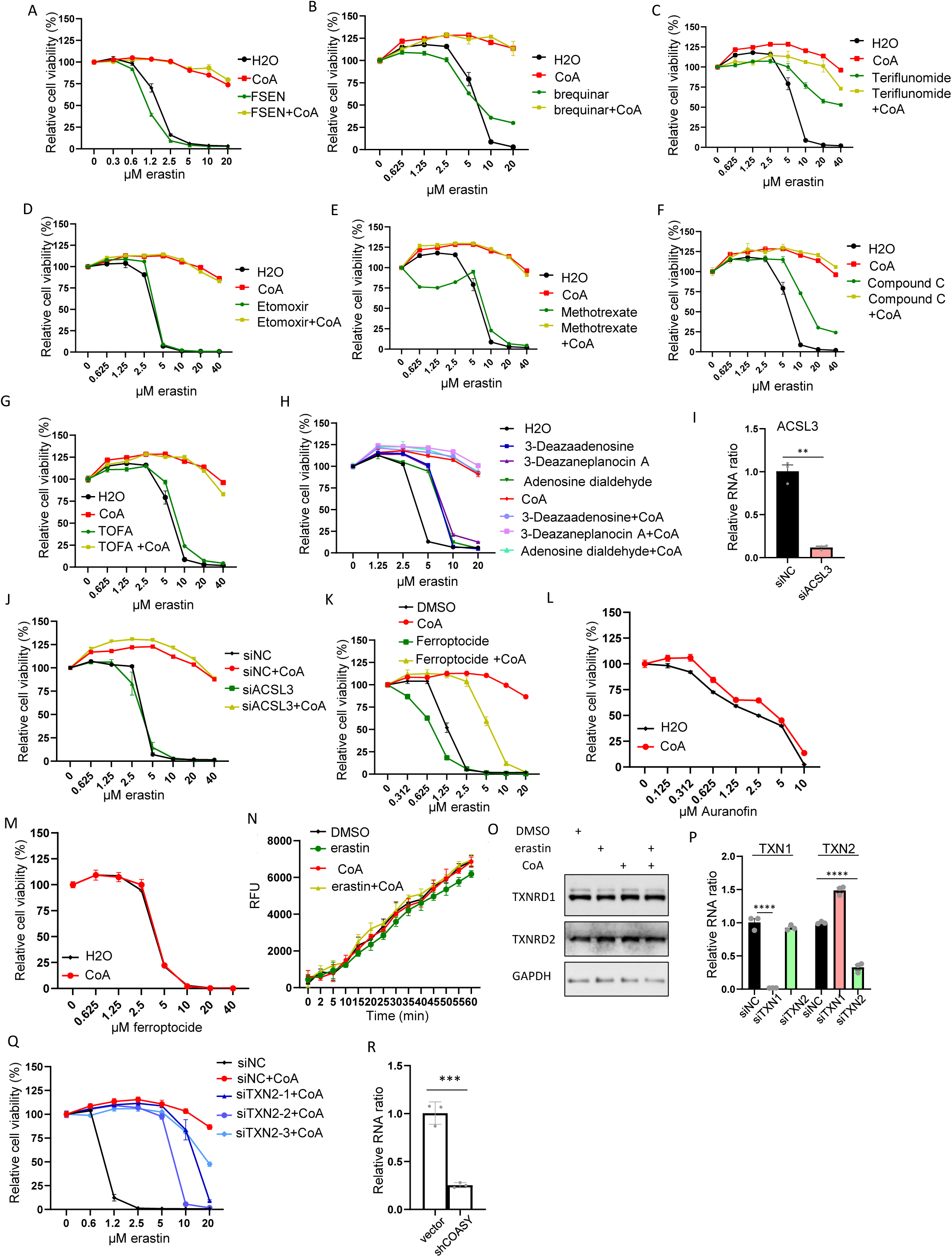
The effects of different genetic and chemical inhibition of candidate targets on the CoA-mediated ferroptosis protection. (**A-K**) Screening for candidate inhibitors or siRNAs that can abolish CoA-mediated ferroptosis inhibition. FSP1 inhibitor (FSEN, 2.5 μM) (**A**), DHODH inhibitor (brequinar, 500 μM) (**B**), DHODH inhibitor (Teriflunomide, 500 μM) (**C**), β-oxidation inhibitor (Etomoxir, 5uM) (**D**), dihydrofolate reductase inhibitor (Methotrexate, 2 μM) (**E**), AMPK inhibitor (compound C, 10 μM) (**F**), ACC inhibitor (TOFA, 25 μM) (**G**), S-adenosyl homocysteine hydrolase inhibitors (3-Deazaadenosine 10 μM, 3-Deazaneplanocin A 10 μM, Adenosine dialdehyde 10 μM) (**H**), siRNA knockdown of ACSL3 (**I-J**), thioredoxin inhibitor (Ferroptocide, 2 μM) (**K**) were treated individually or in combination with CoA (100 μM) upon erastin treatment for 20 hours. The cell viability was quantified by Cell-Titer Glo assay. (**B, C, E, F, G**) were also performed at the same time. Each group has the same control (H2O) and CoA-treated samples. (**L-M**) The CoA failed to rescue auranofin (**L**) or ferroptocide-induced cell death (**M**). HT-1080 cells with CoA (100 μM) were treated with 20 hours of various doses of auranofin (**L**) or ferroptocide (**M**), and cell viability was measured by Cell-Titer Glo assay. (**N**) Individual or combination treatment of erastin (1.25 μM) and CoA (100 μM) for 16 hours did not alter thioredoxin activity in HT-1080 cell lysates. (**O**) Western blots showed that the treatment of erastin (1.25 μM, 16 hours) and CoA (100 μM), either alone or in combination, did not alter TXNRD1 and TXNRD2 protein levels. (**P**) Validation of TXN1 and TXN2 siRNA knockdown in HT-1080 cells. (**Q**) Individual siRNAs targeting three different regions of TXN2 abolished CoA (100 μM)- mediated ferroptosis protection against various doses of erastin for 20 hours. (**R**) Validation of COASY shRNA knockdown by qRT-PCR in HT-1080 cells.

**Supplementary Fig. 3.**
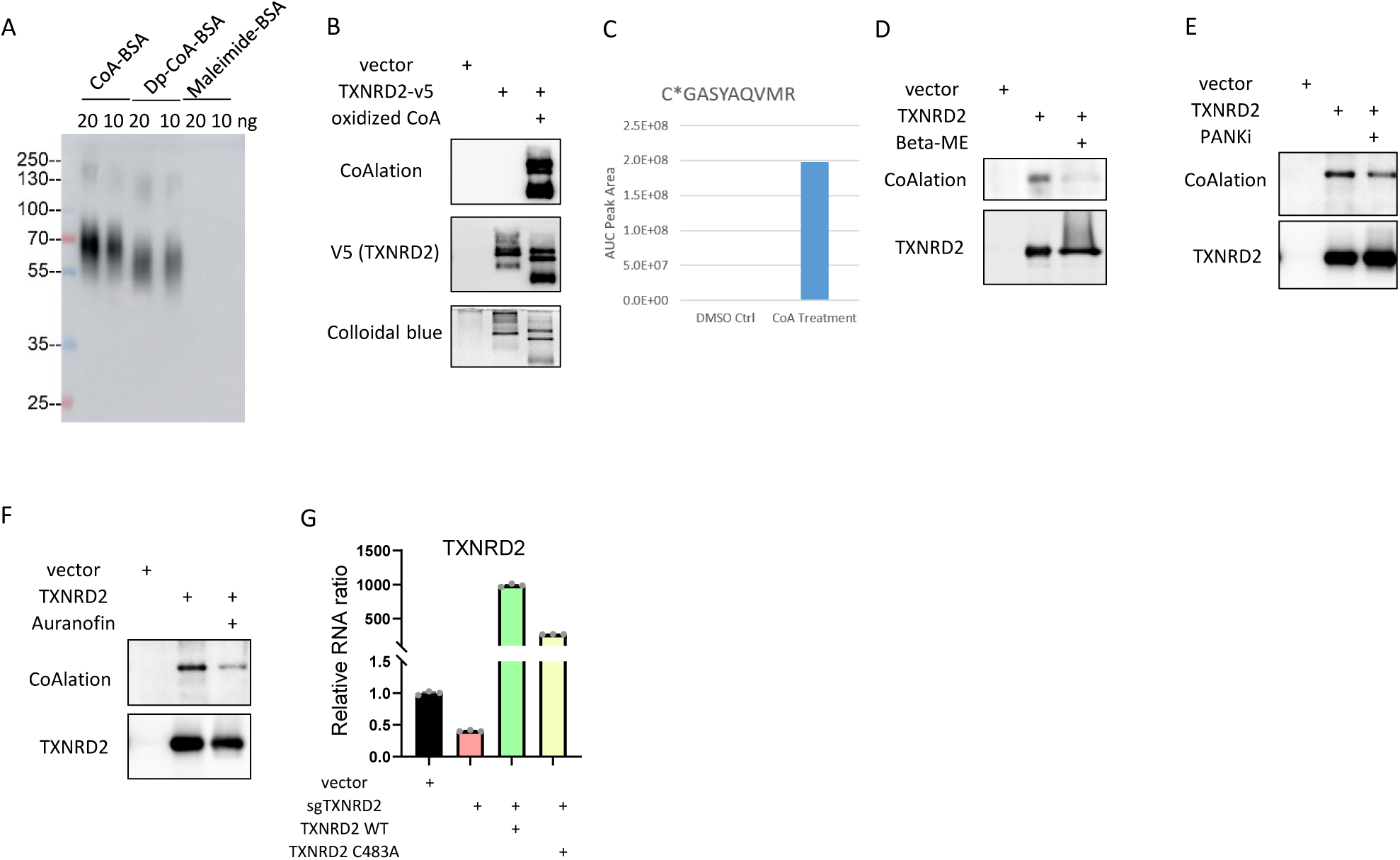
Characterization of the TXNRD2 CoAlation. (**A**) Validation of CoAlation antibody. CoA, 3’-dephospho-CoA (Dp-CoA), or maleimide were conjugated to BSA and resolved on non-reducing SDS PAGE with 20 and 10 ng of protein for Western blots. (**B**) CoAlation on TXNRD2 led to changes in the migration pattern. V5-tag purified TXNRD2 was treated with oxidized CoA for in intro CoAlation, resolved on native PAGE, and stained with Colloidal blue or Western blots for CoAlation. (**C**) Summed EIC peptide intensities from this peptide from each sample. (**D**) Beta-mercaptoethanol (Beta-ME, 100 mM) treatment cleaved the disulfide bond and abolished CoAlation on purified TXNRD2 protein. (**E**) PANKi treatment (5 μM, 20 hours) in HT-1080 cells reduced CoAlation of TXNRD2 protein. (**F**) Auranofin treatment (0.5 μM, 20 hours) in HT-1080 cells reduced CoAlation of TXNRD2 protein. (**G**) Validation of TXNRD2 knockdown by TXNRD2 sgRNA and the over-expression of sgRNA-resistant TXNRD2 wild-type and C483A by Quantitative real-time PCR.

**Supplementary Fig. 4.**
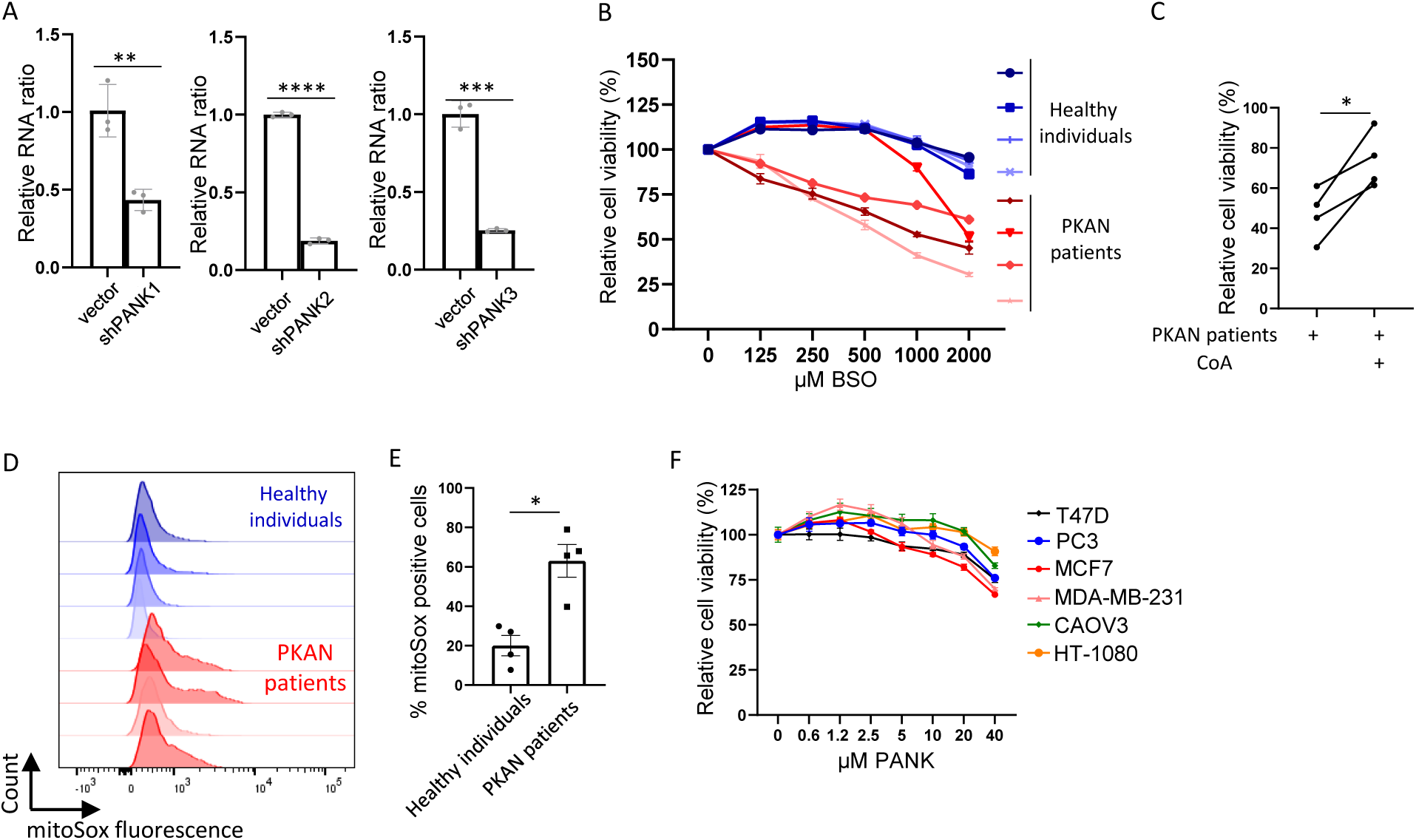
The characterization of PANK as well as PKAN fibroblasts. (**A**) Validation of PANK1, PANK2, and PANK3 shRNA knockdown in HT-1080 cells by RT-PCR. (**B**) The fibroblasts from PKAN patients are sensitive to BSO treatments. The fibroblasts from healthy individuals and PKAN patients were treated with an increasing dose of BSO for 2 days for Cell-Titer Glo assay. (**C**) CoA treatment rescued the PKAN fibroblasts from BSO-induced cell death. Within the same experiment in (**B**), PKAN fibroblasts with BSO were combined with CoA treatment and quantified Cell viability by Cell-Titer Glo assay. (**D**) PKAN fibroblasts, when compared with healthy fibroblasts, showed elevated mitochondrial ROS. Four pairs of fibroblasts from healthy individuals and PKAN patients were stained with the sensor of mitochondrial ROS (mitoSox). (**E**) The quantification of % mitochondrial ROS positive cells in (**D**). (**F)** PANKi treatment did not trigger obvious cell death in the six indicated cancer cell lines.

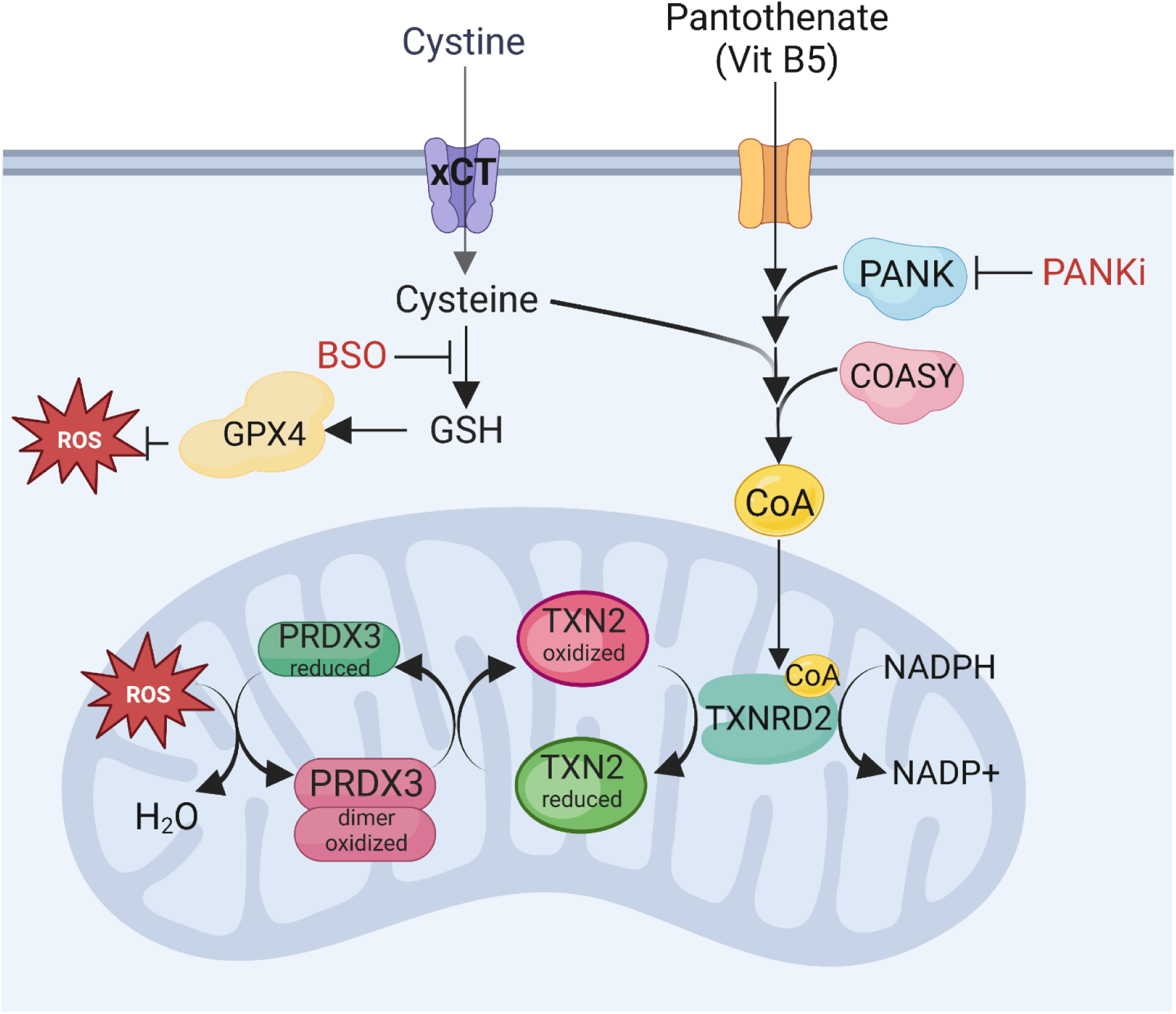

